# Rapid genotyping of viral samples using Illumina short-read sequencing data

**DOI:** 10.1101/2022.03.21.485184

**Authors:** Alex Váradi, Eszter Kaszab, Gábor Kardos, Eszter Prépost, Krisztina Szarka, Levente Laczkó

## Abstract

The most important information about microorganisms might be their accurate genome sequence. Using current Next Generation Sequencing methods, sequencing data can be generated at an unprecedented pace. However, we still lack tools for the automated and accurate reference-based genotyping of viral sequencing reads. This paper presents our pipeline designed to reconstruct the dominant consensus genome of viral samples and analyze their within-host variability. We benchmarked our approach on numerous datasets and showed that the consensus genome of samples could be obtained reliably without further manual data curation. Our pipeline can be a valuable tool for fast identifying viral samples. The pipeline is publicly available on the project’s github page (https://github.com/laczkol/QVG).

## 1. Introduction

The first-hand experience of the severe acute respiratory syndrome coronavirus 2 (SARS-CoV2) pandemic is that effective outbreak management requires fast and strain-level identification of the causative pathogens. The most fundamental information about microorganisms might be their accurately reconstructed genome sequence, which can provide insight into the evolution of pathogens and the clinical outcomes of outbreaks [1]. The application of Next Generation Sequencing (NGS) revolutionized the identification and study of microorganisms by providing an ever-increasing amount of genome sequence data available for data processing and research. Although laboratory instruments are available for numerous research and medical facilities [2], the lack of bioinformatic tools became a bottleneck that hinders high-throughput analysis. Therefore, new widely and openly available bioinformatic tools are needed to keep pace with the increasing speed of data generation and the growing amount of data capable of performing the rapid and accurate analysis of multiple samples sequencing reads.

This paper presents our pipeline designed for the reference-based mass analysis of viral genomes. The pipeline was developed in bash and can be parametrized from the command line. Input samples can be specified using a list of sample file basenames. We paid attention to avoiding the usage of proprietary software to enhance the availability and transparency of the method. We provide a detailed description of the pipeline usage at the project’s github page (https://github.com/laczkol/QVG).

## 2. Description of the pipeline

The pipeline relies on existing tools to characterize samples using NGS data and is designed to readily use the output of any Illumina platform in .fastq format. The method can be applied to both single-end and paired-end sequencing. First, reads are checked for quality and adapter content using fastp 0.20.1 [3] and statistics are exported to .html format. The filtered reads are then aligned to the reference genome sequence using bwa 0.7.17 [4]. Next, duplicate reads are marked with sambamba 0.7.1 [5] and descriptive alignment statistics, including genome coverage, read depth, samtools’ simple statistics (flagstat) and index statistics, are produced with samtools 1.10.2 [6]. Sample files are subset to include only samples covering at least a given proportion of the reference genome (default is 90%). Statistics are plotted using R 3.5 (R Foundation for Statistical Computing, n.d.) and summarized in .pdf files. To capture the polymorphisms of samples two variant calls are performed, both of which use freebayes 1.0.0 [7]. We set freebayes to use the five most probable alleles and annotate variants only with a minimum coverage of five. Base quality scores and mapping quality must have a value larger than 30 to include in variant calling. Alternative alleles with a frequency lower than 20% are excluded from this step. We run freebayes with clumping of haplotypes disabled, hwe priors turned off and use the mapping quality, read placement, strand balance and read position probability instead. To annotate variants of the dominant genome present in a sample ploidy is set to one in the first variant call. Using vcflib 1.0 [8] variants are filtered for a minimum quality of 10 and a ratio of quality / alternate allele observation count of 10 (i.e. each observation is required to have a quality score of at least 10). Then, single-nucleotide polymorphism (SNP) density in 1kbp consecutive windows is extracted from the resulting .vcf files is extracted using vcftools 0.1.16 [9] and visualized using R 3.5 [10]. Vcf statistics as exported by vcfstats 1.0 [8] in plain text files within the output directory. The sequence of the dominant genome is retrieved using vcf2fasta [8]. The filtered reads are aligned to this resulting .fasta file, and regions with a read depth lower than the minimum read depth set for variant calling, are masked out with ‘N’-s using bedtools 2.92.2 [11].

As de novo mutations and/or multiple acquisition sources might introduce genetic heterogeneity of samples [2], [12], a second variant calling step is performed with ploidy unset and assuming pooled sequencing. Low-frequency variants resulting from sequencing error, similar to the first variant call, are filtered out [13]. This step aims to give insight into the population diversity described by allele balance (AB) after filtering the abovementioned variants. The genotypes with their corresponding AB are saved to plain text files using bcftools 1.9 [14] and visualized using R 3.5 [10].

This way, running the pipeline exports the dominant viral genome of samples and provides insight into the intra-host diversity. With the usage of GNU parallel [15], tasks are run in parallel to decrease computation time. Using Liftoff 1.6.3 [16], annotations of the reference genome (in gff3 format) can be transferred to the consensus sequences output by QVG.

### 2.1. Benchmarking

The pipeline presented in this study was tested on different operating systems, namely, Ubuntu Server 20.04, Linux Mint 20.2, Debian 10.1, and 11.0, but it can run using any UNIX-like operation system with dependencies installed correctly. Requirements of the pipeline were installed using the conda package manager as specified in the yaml configuration file uploaded to the github repository of the pipeline (https://github.com/laczkol/QVG). The accuracy and performance of the pipeline were tested on multiple datasets that we describe below. In the first run, samples of the given dataset (Table 1) were analyzed simultaneously using six CPU cores; then, to assess the correlation of running time, coverage, and the number of reads supplied for the run, samples were genotyped one by one using one CPU core.

**Table 1.**
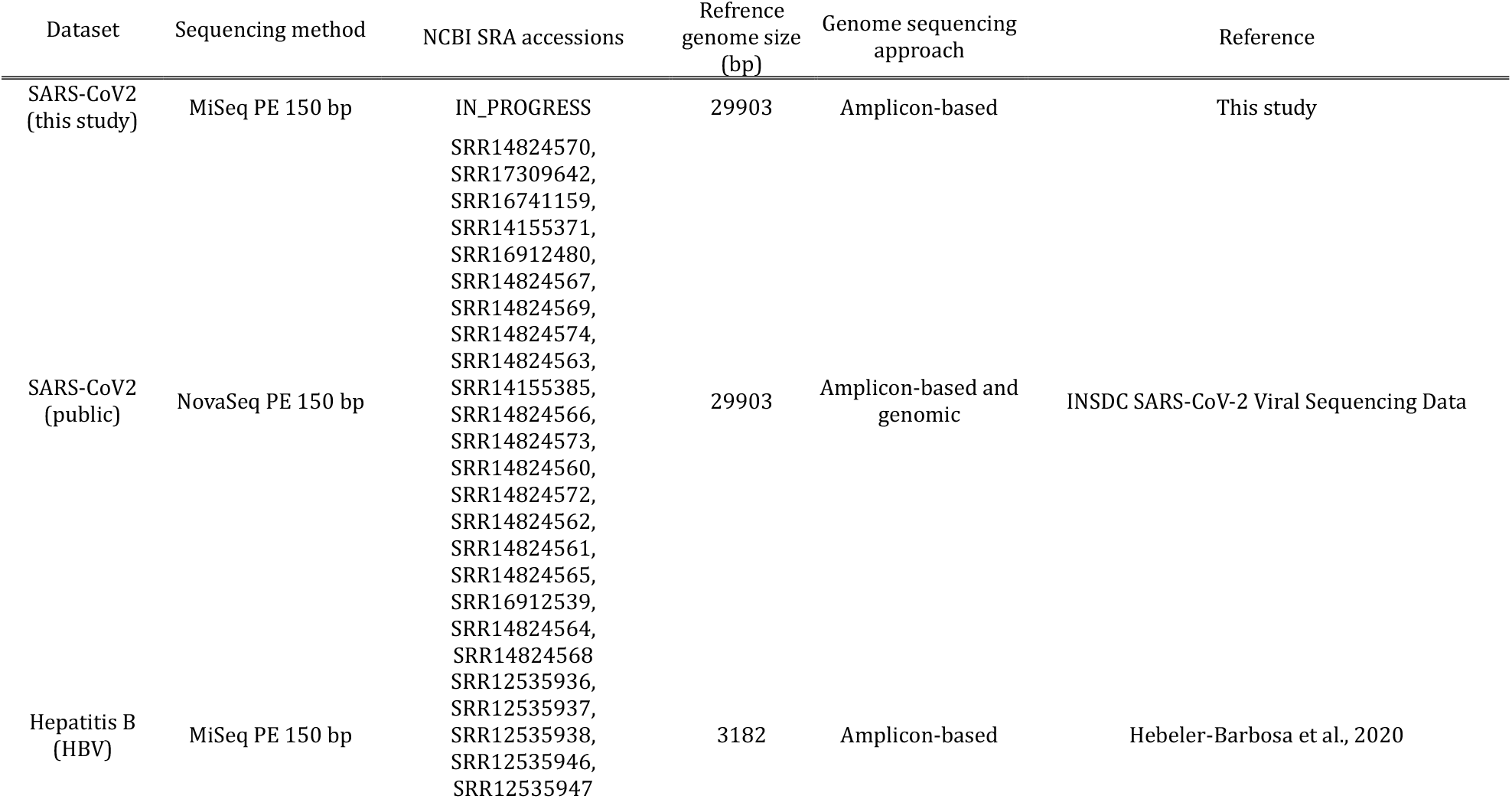
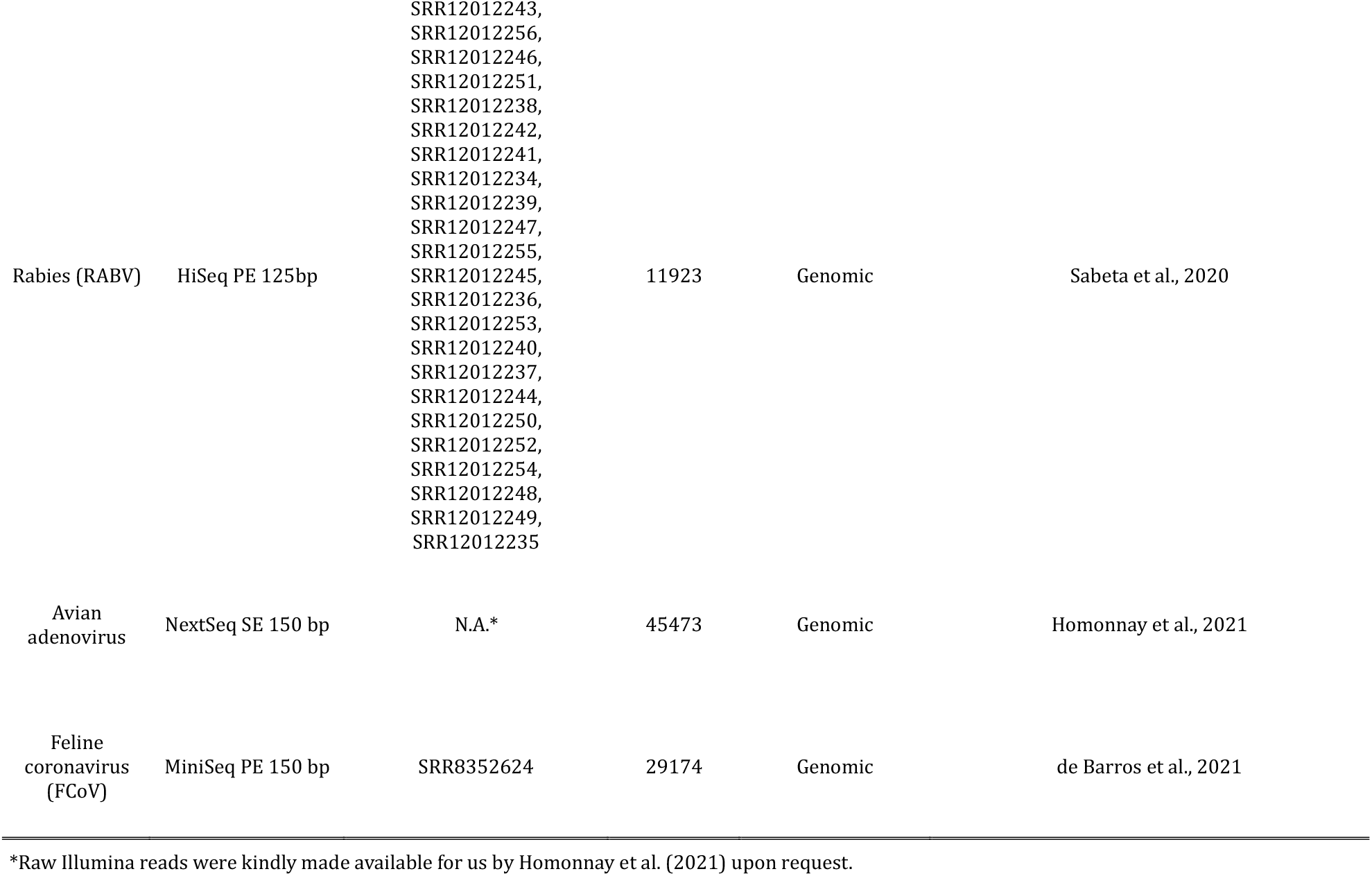
Summary of datasets used in this study.

We tested the performance of our pipeline by comparing the results obtained by Quick Viral Genome Genotyper (QVG) against the output of Geneious Prime 2021.2.2. Owing to its ease of use, Geneious is one of the most widely used cross-platform commercial software to carry out reference-based genotyping of samples. For this comparison, we used 20 SARS-CoV-2 positive nasopharyngeal samples (Table 1) (New Coronavirus Nucleic Acid Detection Kit (Perkin Elmer, Waltham, MA, USA); samples with <30 threshold cycle were chosen) to sequence the virus genome. RNA was extracted using Viral DNA/RNA extraction kit and Automated Nucleic Acid Extraction System-32 (BioTeke Corporation, Beijing, China) then libraries were prepared with NEXTFLEX® Variant-Seq™ SARS-CoV-2 Kit (For Illumina® Platforms) (Perkin Elmer, Waltham, MA, USA). The libraries were processed in an Illumina MiSeq platform using a MiSeq Reagent Kit v3 (Illumina, San Diego, CA, USA) following the manufacturers’ instructions. As a reference sequence for this experiment, we used the genome of the SARS-CoV2 isolate Wuhan-Hu-1 (MN908947.3). In Geneious, after removing duplicate reads, reads were mapped to the reference genome using the Geneious mapper with the default sensitivity (Medium Sensitivity/Fast). Before mapping, sequences were trimmed the same way as in the QVG pipeline. As a final step, we checked the alignment of reads for obvious genotyping errors and corrected the consensus genome manually. This procedure took ~10-15 minutes per sample. Geneious was run on a computer with an Intel Core i7-11700K 3.60GHz CPU running Windows 10 64-bit. Using the parameters described above, the genome sequences obtained by both approaches were submitted to the Pangolin web server (https://cov-lineages.org/resources/pangolin.html) to assign each sample to its corresponding lineage.

In addition, we collected and re-analyzed publicly available SARS-CoV-2 sequencing reads with known identity (Table 1). These raw sequencing data were either produced by amplicon-based sequencing (lineages Alpha, Beta, Gamma, Epsilon, Eta) or a transcriptomic sequencing approach (lineage Omicron). Publicly available samples were genotyped, relying on the same reference genome and parameter values we used for our newly generated sequencing data. Re-analyzed consensus genome sequences were submitted to the Pangolin webserver (https://cov-lineages.org/resources/pangolin.html), then we compared the assigned lineage to the originally reported one (see Table 1. for accession numbers).

Another amplicon sequencing-based dataset used for the benchmarking was the dataset presented by [17]. Raw reads of 17 Hepatitis B (HBV) virus samples were supplied to our pipeline. Samples were genotyped using the read alignment to the reference genome of the Hepatitis B virus (strain ayw) (NC_003977). The consensus genome sequences were submitted to the Genome Detective’s HBV phylogenetic typing tool (https://www.genomedetective.com/app/typingtool/hbv/ [18]). This tool not only reports the most probable lineage assigned to samples but conducts a recombination analysis using bootscan [19]. We compared the genotypes assigned by Genome Detective with the originally reported lineage by Hebeler-Barbosa et al. (2020) [17].

To demonstrate that our approach can process not only AmpliSeq datasets, we run the Rabies virus (RABV) dataset presented by Sabeta et al. (2020) [20] through our pipeline. Sequencing reads of this dataset were obtained after the depletion of host DNA and RNA [20]. To genotype the samples of this dataset, we used the genome of Rabies virus (isolate 20034) (KT336433). We checked the identity of samples by submitting the consensus genome sequences to the RABV-GLUE identification tool (http://rabv-glue.cvr.gla.ac.uk/), then compared the most probable lineage uncovered by this tool with the identity of the originally reported lineage. Furthermore, we re-analyzed the feline coronavirus (FCoV) sequence data of de Barros et al. (2019) [21] and the avian adenovirus sequencing reads of Homonnay et al. (2021) [22], latter of which was the only single-end read sequencing dataset included in the benchmarking. For the FCoV dataset, we reduced the minimum read depth required for variant calling to three, as this sample showed the lowest vertical coverage after aligning the reads to the reference genome of feline coronavirus (isolate UG-FH8) (KX722529) also used by de Barros et al. (2019) [21]. The genotyping of the adenovirus sample used the reference genome of the fowl aviadenovirus B strain (40440-M/2015) (MG953201). Since no subtyping tool exists for the latter two viral species, we used blastn to match the consensus genome sequence against the NCBI nucleotide collection database; then the retrieved highest-scoring pairs were subject of phylogenetic reconstruction with fasttree [23] and pairwise distance matrix calculation using the proportion of different sites between samples (‘raw’ distance) as implemented in the R package pegas [24].

## 3. Results and Discussion

We could obtain a good quality reference in all runs presented here. The most important factor to influence the total running time (including the quality filtering, read alignment, and variant calling) appeared to be the number of reads supplied to the pipeline, regardless of the sequencing approach (Figure 1.). The mean read depth of the samples affected the running time to a much lesser extent than the total number of reads (including those that did not align to the reference genome). The running time varied considerably; the SARS-CoV2 sample S5 generated for this study could be genotyped under 18 seconds, whereas the analysis of the SARS-CoV2 sample SRR14824569 needed the most time to finish, more than 43 minutes. Both extremities of running time used an amplicon-based approach to obtain sequencing reads. The genotyping of the samples relying on a genomic approach could be run in a similar time span. The only SARS-CoV2 sample relying on the transcriptomic approach (SRR17309642) was analyzed under two minutes, whereas the RABV sample SRR12012239 could be processed in 39 minutes (Figure 1.). The total running time could be decreased proportionally by using more CPU cores.

**Figure 1.**
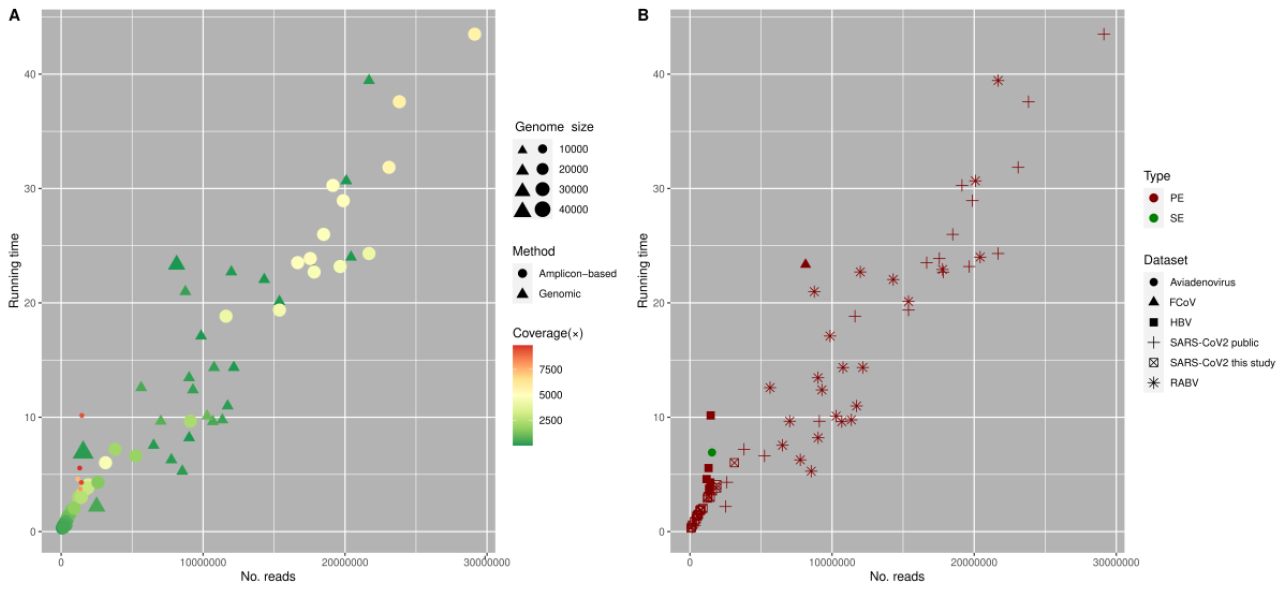
Time required in minutes to run the whole pipeline as reported by the ‘time’ utility using one CPU core as a function of genome coverage. This analysis was run on a commercial laptop with an Intel i7-4910MQ processor. Using more threads decreased the running time proportionally. On the left plot (A) the size of symbols is proportional to the reference genome size. Different symbols indicate the approach used for genome sequencing. The “genomic” approach includes whole genome, metagenomic and transcriptome sequencing. The symbols color represent the number of reads supplied for the pipeline including those that could not be aligned to the reference genome. On the right panel (B) the symbols color show the type of the sequencing run and different symbols indicate the samples corresponding dataset (Table 1.).

Sequencing reads of the SARS-CoV2 dataset generated for this study covered 94.7—99.9 % of the reference genome (Table S1.). The SNP density of all 20 samples appeared to be roughly equal across the genome, except at ORF8, in line with the findings of Flower et al. (2021) [25], and in the gene encoding the spike protein (S) that is known to harbor several mutations in the lineage AY.4 (Figure 2). Polymorphism within samples indicating more than one probable allele (Figure 3.) could be found in all samples, but the same polymorphic site rarely showed an AB > 0 in more than one sample. Generally, 1–5 sites showed within-host variability. The only exceptions were two transitions at positions 21,987 and 24,410 found in 17 and 12 isolates, respectively. These are known but not characteristic mutations of the lineage identified by Pangolin. Submitting the alternative alleles to Pangolin did not change the result of the lineage assignment. The Pangolin lineage assignment using the consensus genome obtained by QVG and Geneious showed identical results and very similar support values, except for the sample S15. This sample using QVG could be assigned to the lineage AY.42, whereas using Geneious, it could be identified as AY.43. This discordance could be linked to this sample’s relatively lower genome coverage (Table S1.). Statistical support values were not unequivocally better for either pipeline (Table 2.).

**Figure 2.**
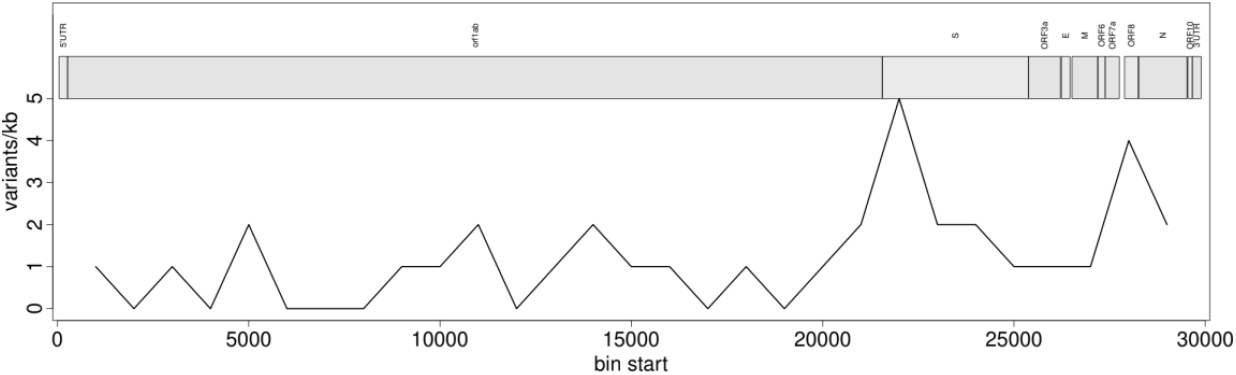
Example of the SNP density of sample S11 across the reference genome. The x-axis shows the genomic position, whereas the y-axis represents the number of SNPs within sliding windows.

**Figure 3.**
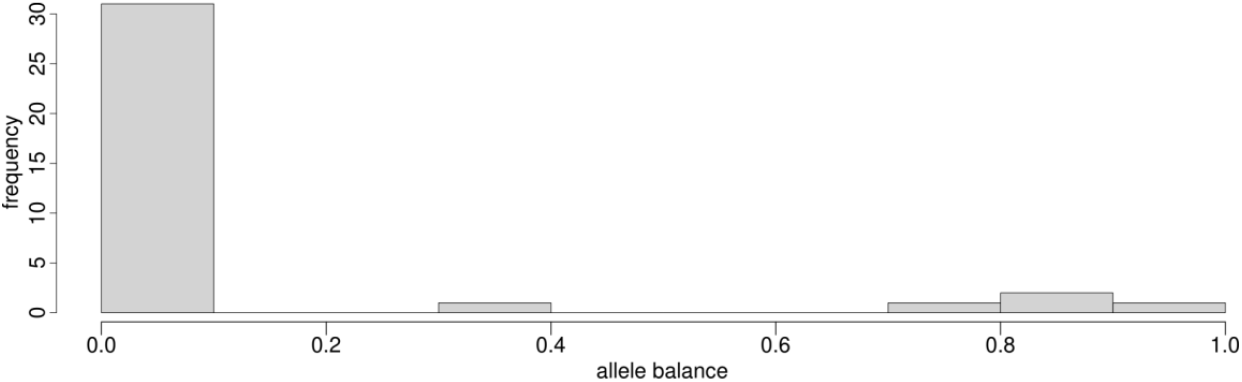
Example of AB distribution (sample S11) visualized as a histogram. An AB value different from 0 suggests multiple probable alleles at a given site.

**Table 2.** Comparison of pipelines used in this study by the lineage assignment and support values as output by Pangolin. The only sample assigned differently after genotyping by the two compared pipelines is given in bold.

| Sequence name | QVG |  |  |  |  |  | Geneious |  |  |  |  |  |
| --- | --- | --- | --- | --- | --- | --- | --- | --- | --- | --- | --- | --- |
|  | Lineage | Conflict | Am-biguity score | Scorpio call | Scorpio support | Scorpio conflict | Lineage | Conflict | Am-biguity score | Scorpio call | Scorpio support | Scorpio conflict |
| S2 | AY.4 | 0 | 1 | Delta (AY.4-like) | 0.91 | 0.06 | AY.4 | 0 | 1.00 | Delta (AY.4-like) | 0.91 | 0.03 |
| S3 | AY.46.6 | 0 | 0.96 | Delta (B.1.617.2-like) | 0.85 | 0.15 | AY.46.6 | 0 | 0.97 | Delta (B.1.617.2-like) | 0.92 | 0.08 |
| S4 | AY.46 | 0 | 0.99 | Delta (B.1.617.2-like) | 0.92 | 0.08 | AY.39 | 0 | 0.99 | Delta (B.1.617.2-like) | 0.85 | 0.08 |
| S5 | AY.43 | 0 | 0.99 | Delta (B.1.617.2-like) | 0.92 | 0.08 | AY.43 | 0 | 0.99 | Delta (B.1.617.2-like) | 0.92 | 0.08 |
| S8 | AY.4 | 0 | 1 | Delta (AY.4-like) | 0.91 | 0.06 | AY.4 | 0 | 1 | Delta (AY.4-like) | 0.94 | 0.03 |
| S9 | AY.43 | 0 | 1 | Delta (B.1.617.2-like) | 0.92 | 0.08 | AY.43 | 0 | 1 | Delta (B.1.617.2-like) | 0.92 | 0.08 |
| S10 | AY.43 | 0 | 1 | Delta (B.1.617.2-like) | 0.92 | 0.08 | AY.43 | 0 | 1 | Delta (B.1.617.2-like) | 0.92 | 0.08 |
| S11 | AY.9.2 | 0 | 1 | Delta (B.1.617.2-like) | 0.92 | 0.08 | AY.9.2 | 0 | 1 | Delta (B.1.617.2-like) | 1 | 0 |
| S12 | AY.9.1 | 0 | 1 | Delta (B.1.617.2-like) | 0.92 | 0.08 | AY.9.1 | 0 | 1 | Delta (B.1.617.2-like) | 1 | 0 |
| S13 | AY.43 | 0 | 1 | Delta (B.1.617.2-like) | 0.92 | 0.08 | AY.43 | 0 | 1 | Delta (B.1.617.2-like) | 0.92 | 0.08 |
| S14 | AY.43 | 0 | 1 | Delta (B.1.617.2-like) | 0.92 | 0.08 | AY.43 | 0 | 1 | Delta (B.1.617.2-like) | 0.85 | 0.15 |
| **S15** | **AY.42** | **0** | **0.94** | **Delta (B.1.617.2-like)** | **0.69** | **0.15** | **AY.43** | **0** | **0.96** | **Delta (B.1.617.2-like)** | **0.85** | **0** |
| S16 | AY.3 | 0 | 1 | Delta (B.1.617.2-like) | 0.92 | 0.08 | AY.3 | 0 | 1 | Delta (B.1.617.2-like) | 0.92 | 0.08 |
| S17 | AY.43 | 0 | 1 | Delta (B.1.617.2-like) | 0.92 | 0.08 | AY.43 | 0 | 1 | Delta (B.1.617.2-like) | 0.92 | 0.08 |
| S18 | AY.122 | 0 | 1 | Delta (B.1.617.2-like) | 0.92 | 0.08 | AY.122 | 0 | 1 | Delta (B.1.617.2-like) | 0.92 | 0.08 |
| S19 | AY.46.6 | 0 | 1 | Delta (B.1.617.2-like) | 1 | 0 | AY.46.6 | 0 | 1 | Delta (B.1.617.2-like) | 1 | 0 |
| S20 | AY.43 | 0 | 1 | Delta (B.1.617.2-like) | 0.92 | 0.08 | AY.43 | 0 | 1 | Delta (B.1.617.2-like) | 0.92 | 0.08 |
| S22 | AY.43 | 0 | 1 | Delta (B.1.617.2-like) | 1 | 0 | AY.43 | 0 | 1 | Delta (B.1.617.2-like) | 1 | 0 |
| S23 | AY.122 | 0 | 0.99 | Delta (B.1.617.2-like) | 0.92 | 0.08 | AY.122 | 0 | 0.99 | Delta (B.1.617.2-like) | 0.92 | 0.08 |
| S24 | AY.122 | 0 | 1 | Delta (B.1.617.2-like) | 1 | 0 | AY.122 | 0 | 1 | Delta (B.1.617.2-like) | 1 | 0 |

The publicly available SARS-CoV2 sequencing data showed similar results. The genome coverage varied between 97.1—100 % (Table S2). Similar to the dataset generated for this study, SNPs showed the highest density at the spike protein and ORF8. Only two samples did not show signs of within-host diversity (SRR16912539, SRR16912480). Other samples showed 1—15 polymorphisms with an AB > 0, of which SNPs at positions 28,270 could be found in 11 samples, whereas such polymorphisms at positions 28,095 and 29,870 were found in 4— 4 samples. The identities of the consensus genomes always matched with the already published identification (Table 3.). Only sample SRR14824567 was classified as a different lineage (B.1.637) than the original lineage (B.1.526.1)., but later this B.1.526.1 was designated to B.1.637 in Pangolin. Despite these samples having a various number of reads, mean read depth, and being sequenced using different approaches (Table 1., Table S2.), our pipeline outputs good quality consensus genomes.

**Table 3.** Comparison of originally reported lineages, and lineages identified by Pangolin after genotyping publicly available sequencing reads of SARS-CoV2 with our pipeline.

| Sequence name | Lineage | Conflict | Ambiguity score | Scorpio call | Scorpio support | Scorpio conflict | Originally reported lineage |
| --- | --- | --- | --- | --- | --- | --- | --- |
| SRR14155371 | B.1.1.7 | 0 | 0.98 | Alpha (B.1.1.7-like) | 0.96 | 0.04 | B.1.1.7 |
| SRR14155385 | B.1.1.7 | 0 | 1.0 | Alpha (B.1.1.7-like) | 0.96 | 0.04 | B.1.1.7 |
| SRR14824560 | B.1.1.7 | 0 | 0.98 | Alpha (B.1.1.7-like) | 0.96 | 0.04 | B.1.1.7 |
| SRR14824561 | B.1.1.7 | 0 | 0.98 | Alpha (B.1.1.7-like) | 0.96 | 0.04 | B.1.1.7 |
| SRR14824562 | B.1.429 | 0 | 1.0 | Epsilon (B.1.429-like) | 1.0 | 0 | B.1.429 |
| SRR14824563 | P.1 | 0 | 1.0 | Gamma (P.1-like) | 0.87 | 0 | P.1 |
| SRR14824564 | B.1.1.7 | 0 | 1.0 | Alpha (B.1.1.7-like) | 0.91 | 0.04 | B.1.1.7 |
| SRR14824565 | B.1.1.7 | 0 | 1.0 | Alpha (B.1.1.7-like) | 0.96 | 0.04 | B.1.1.7 |
| SRR14824566 | P.1 | 0 | 1.0 | Gamma (P.1-like) | 0.87 | 0 | P.1 |
| SRR14824567 | B.1.637 | 0 | 1.0 |  |  |  | B.1.526.1 |
| SRR14824568 | B.1.1.7 | 0 | 1.0 | Alpha (B.1.1.7-like) | 0.95 | 0.04 | B.1.1.7 |
| SRR14824569 | B.1.1.7 | 0 | 1.0 | Alpha (B.1.1.7-like) | 0.95 | 0.04 | B.1.1.7 |
| SRR14824570 | B.1.1.7 | 0 | 1.0 | Alpha (B.1.1.7-like) | 0.95 | 0.04 | B.1.1.7 |
| SRR14824572 | B.1.525 | 0 | 0.98 | Eta (B.1.525-like) | 1.00 | 0 | B.1.525 |
| SRR14824573 | B.1.1.7 | 0 | 1.0 | Alpha (B.1.1.7-like) | 0.96 | 0.04 | B.1.1.7 |
| SRR14824574 | B.1.1.7 | 0 | 1.0 | Alpha (B.1.1.7-like) | 0.96 | 0.04 | B.1.1.7 |
| SRR16741159 | B.1.351 | 0 | 0.98 | Beta (B.1.351-like) | 0.78 | 0.14 | B.1.351 |
| SRR16912480 | P.1 | 0 | 1.0 | Gamma (P.1-like) | 0.87 | 0 | P.1 |
| SRR16912539 | P.1 | 0 | 1.0 | Gamma (P.1-like) | 0.87 | 0 | P.1 |
| SRR17309642 | BA.1 | 0 | 1.0 | Omicron (BA.1-like) | 0.91 | 0 | B.1.1.529/Omicron |

The genome coverage of the HBV dataset showed a higher variability (65.7—100 %). SNP density in 1,000 bp windows peaked at 84 (sample SRR12535947) and generally showed a decreasing trend towards the end position of the reference genome. Samples had 1—28 polymorphic sites with more than one probable alleles. The same ‘non-haploid’ (i.e. multiple probable alleles could be observed) position could be observed in a maximum of two samples. Genome Detective could correctly assign genomes into HBV subtypes, except for one sample (Table 4.). The bootscan analysis (Figure 4.) confirmed that the dominant genome, which could not be equivocally assigned to any lineage, can be a recombinant of strains A and D. Recombination is not unprecedented for HBV [26],[27],[28] and can play an important role in the evolution of HBV genotypes [26].

**Table 4.** Short result of the phylogenetic type classification of HBV samples by Genome Detective.

| Name | Length | Begin | End | Species | Type | Type support | Original subtype |
| --- | --- | --- | --- | --- | --- | --- | --- |
| SRR12535936 | 3182 | 1 | 3182 | Hepatitis B virus | subtype D | 100.0 | subtype D |
| SRR12535937 | 3179 | 4 | 3182 | Hepatitis B virus | subtype D | 100.0 | subtype D |
| SRR12535938 | 3182 | 1 | 3182 | Hepatitis B virus | subtype D | 100.0 | subtype D |
| SRR12535946 | 3182 | 1 | 3182 | Hepatitis B virus | subtype D | 100.0 | subtype D |
| SRR12535947 | 3179 | 1 | 3182 | Hepatitis B virus | Could not assign |  | subtype A |

**Table 5.**
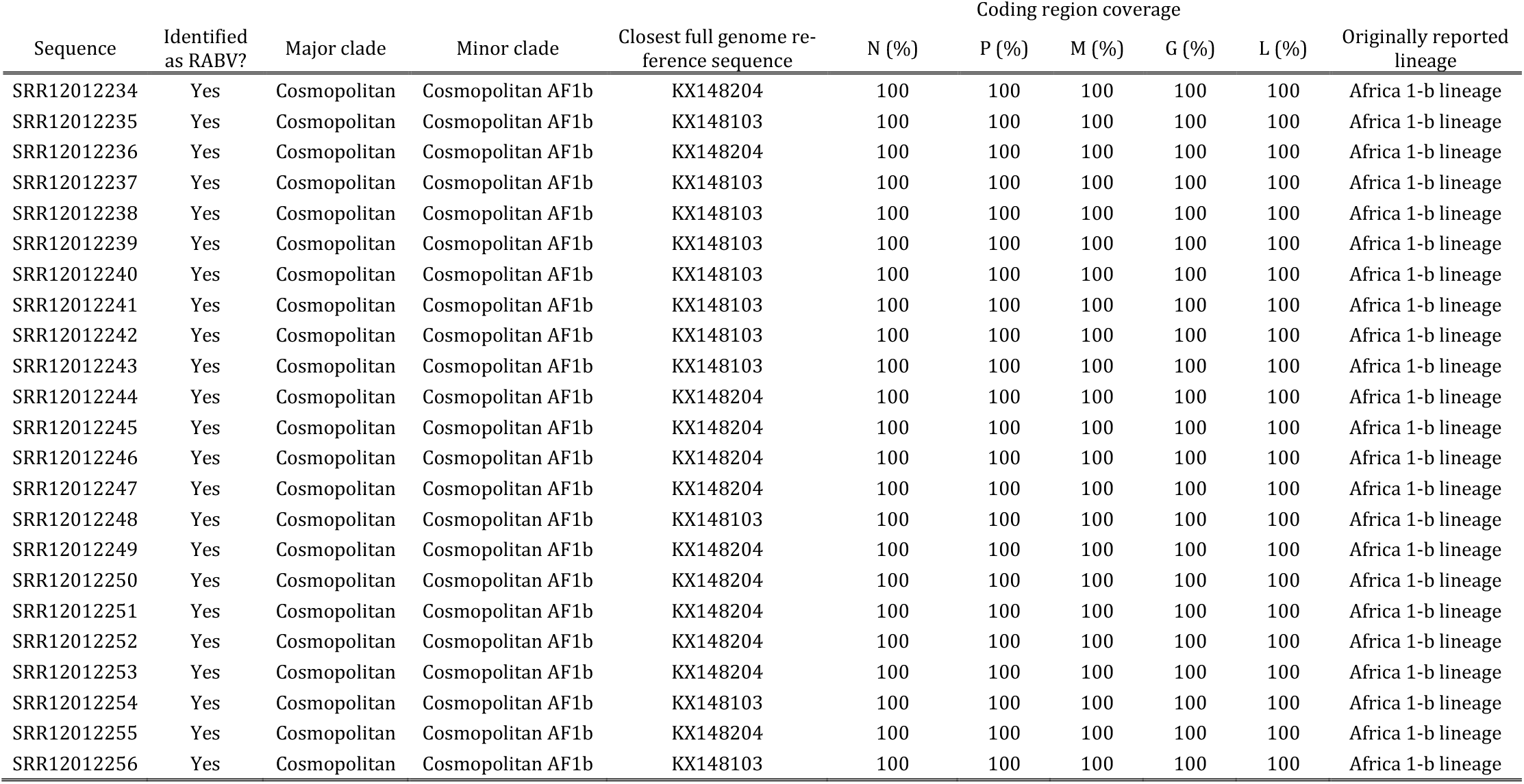
Result of classification of the RABV datasets samples returned by RABV-GLUE.

**Figure 4.**
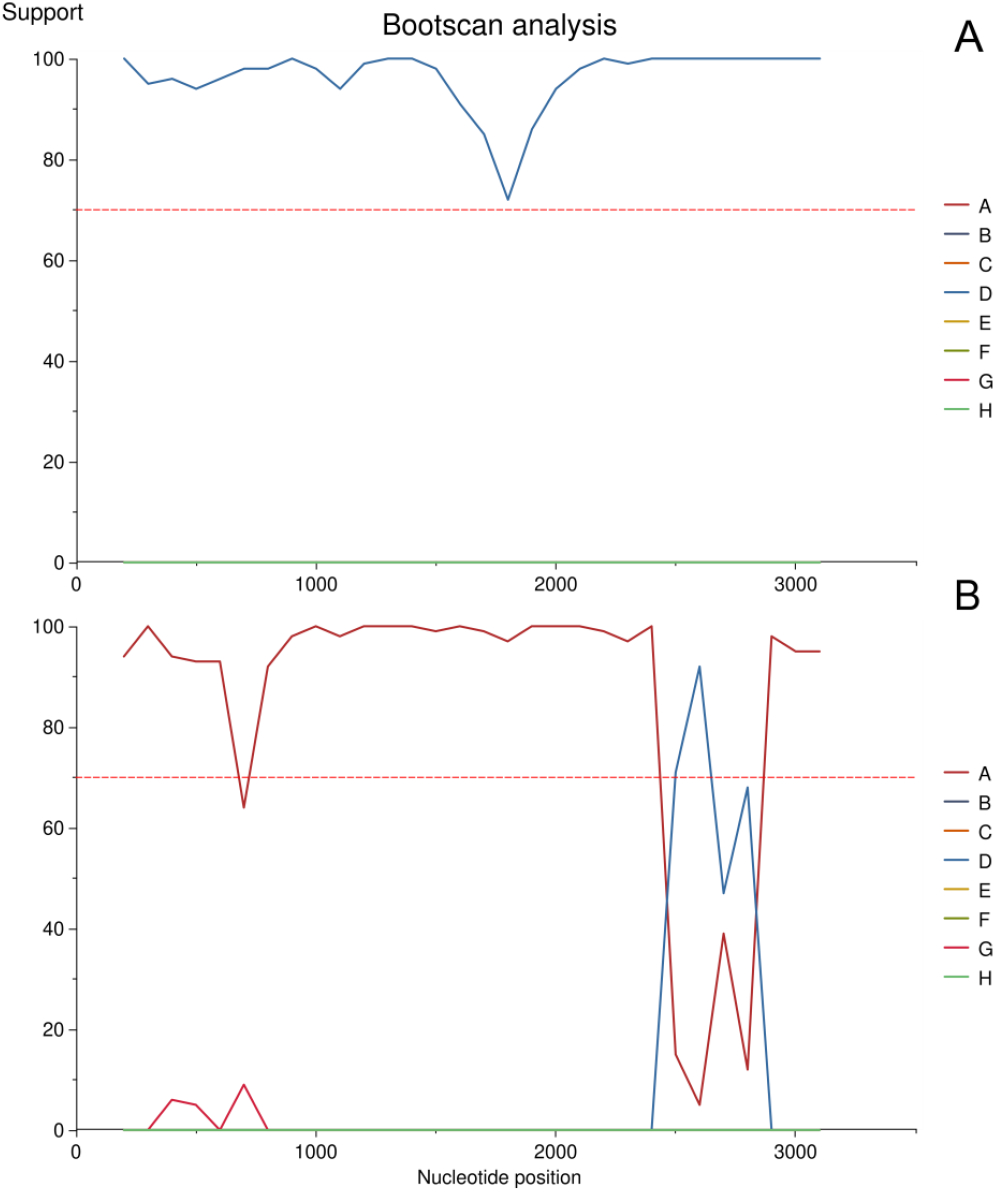
Example of an unequivocally identified HBV sample (A) (SRR12535946) and a recombinant sample (B) (SRR12535947) as output by Genome Detective using Bootscan.

The genome coverage of RABV samples appeared to be at least 98.76 %. Since the genomic approach applied to obtain the sequencing reads of this dataset does not strictly rely on species-specific PCR amplicons, the horizontal coverage (Table S4.) was lower than for previously described datasets. SNP density of the dataset varied between 11—37 and showed a roughly uniform distribution within samples. Seven out of 23 samples showed no variants with an AB > 0 (SRR12012247, SRR12012251, SRR12012242, SRR12012245, SRR12012250, SRR12012240, SRR12012237, SRR12012254). The remaining samples had 1—14 ‘non-haploid’ sites, and the same such site could be observed in a maximum of two samples. RABV-GLUE identified the samples as the cosmopolitan AF1b lineage, agreeing with the originally reported classification of Sabeta et al. (2020).

Sequencing reads of the FCoV sample covered more than 94 % of the reference genome but showed the lowest coverage of all samples analyzed in this study (Table S5.). The SNP density peaked at 73 and showed multiple highly polymorphic islands along the reference genome. In total, 72 out of 1,411 polymorphic sites showed within-host diversity based on AB values. The phylogenetic reconstruction correctly placed the consensus genome output by QVG closest to the publicly available genome (Figure 5A) of the same sample. However, a relatively higher genetic distance could be observed between these two sequences (Figure 5A and B). We link this phenomenon to the relatively low read depth of the sequencing reads (mean = 4.87), which can decrease reference-based genotyping accuracy. The low read depth can be a limitation of the approach presented here and any reference-based genotyping method.

**Figure 5.**
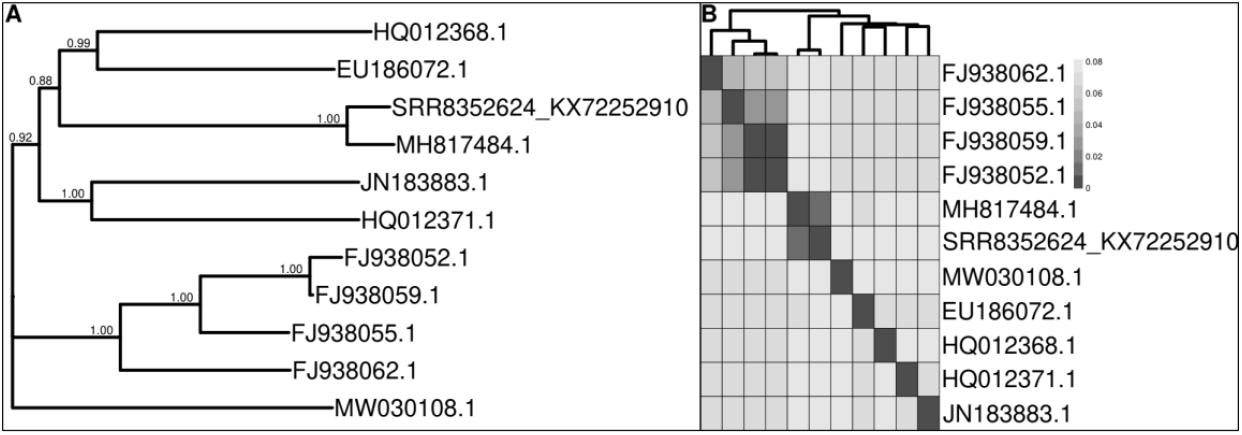
Phylogenetic tree reconstructed for the best 10 BLAST hits using the consensus FCoV genome obtained using our pipeline (A) and pairwise sequence similarity shown on a heatmap of these sequences using raw distances (B). The sample name SRR8352624_KX72252910 represents the sequencing reads genotyped with our pipeline relying on alignments to the reference genome KX722529.1 and MH817484 shows the position of the publicly available reference genome of feline coronavirus strain FCoV-SB22 (de Barros et al., 2019).

The obtained consensus genome of the adenovirus sample covered 99.4 % of the reference genome. The SNP density appeared to be higher in the pVI, ORF22, and ORF17—ORF19A genes relative to the rest of the genome. In total, we observed seven sites with an AB > 0. The phylogenetic reconstruction clustered the consensus genome genotyped here and the publicly available genome of the same sample (Figure 6A), agreeing with the clustering based on pairwise genetic distances. We only observed indel mutations between the two mentioned sequences that could be linked to the automatic masking of low-depth genomic regions (read depth < 5). This low divergence of the reference points out the accuracy of the presented pipeline.

**Figure 6.**
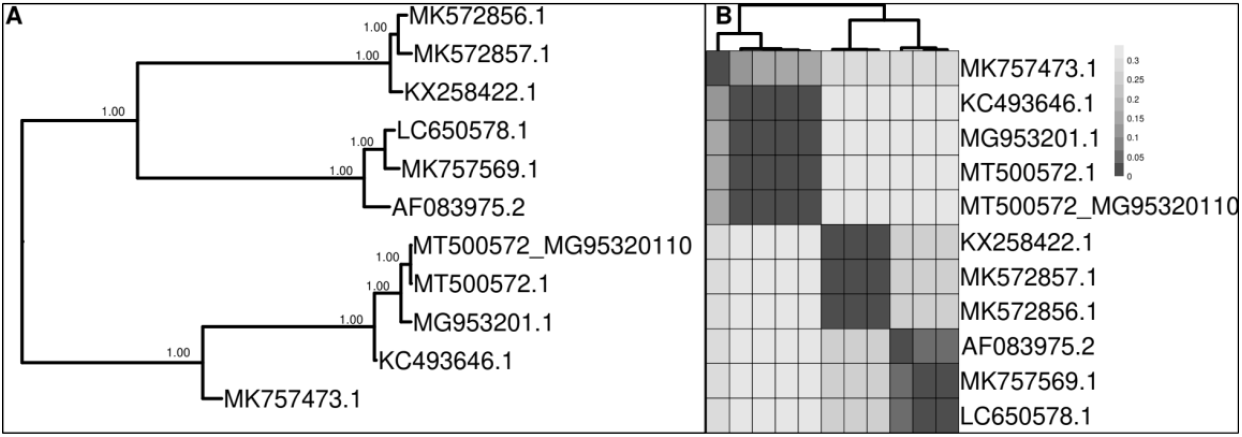
Phylogenetic tree reconstructed for the best 10 BLAST hits using the consensus avian Adenovirus genome obtained using our pipeline (A) and pairwise sequence similarity shown on a heatmap of these sequences using raw distances (B). The sample name MT500572_MG95320110 represents the sequencing reads genotyped with our pipeline relying on alignments to the reference genome MG953201.1 and MT500572.1 shows the position of the publicly available reference genome of the avian adenovirus isolate D2453/1/10-12/13/UA (Homonnay et al., 2021).

This work reports a pipeline capable of rapid and automated analysis of viral genomes obtained by NGS. Unlike proprietary software solutions, this pipeline relies on freely available, open-source bioinformatic software. Using parallel execution of tasks, we could obtain consensus genomes of the SARS-CoV2 dataset generated for this study without the need for laborious manual data curation required by Geneious and with similar accuracy. Our pipeline generated good quality consensus genomes using its default settings in most cases, with the FCoV sample as the only exception. Moreover, we could also investigate the intra-host diversity of samples using the allele balance values. The occurrence of the same variable sites sharing more probable identical alternative alleles within datasets showed that these ‘non-haploid’ polymorphisms are probably existing mutations originating from multiple acquisitions of different strains.

We genotyped already known genomes of four viral species. Some of these viruses are highly variable (HBV, SARS CoV-2) and can pose dangers to humans and domestic or wild animals (RABV, FcoV); thus, it can be important to identify them and to track their molecular evolution. All samples genotyped by our pipeline were correctly identified by classification tools, except the HBV samples that appeared to be a recombinant genome. Our results demonstrate that QVG can handle a wide range of Illumina sequencing platforms (NextSeq, MiniSeq, MiSeq, HiSeq 2500, NovaSeq 6000), different genome sizes (3182 – 29903 base), a broad range of short read lengths (76 – 150bp) and a good accuracy could be achieved regardless of these parameters. However, care should be taken to set the correct parameters if the reference genome coverage or the mean read depth is relatively lower. The only inconsistency between genotyping approaches (S15 of the SARS-CoV2 dataset) and an inflated genetic distance (FCoV) could be linked to these issues.

The obvious shortcoming of this method is that the characterization of samples relies on a closely related reference genome, which, if not yet available, should be assembled first using e.g. VirusTAP [29] or V-ASAP [2]. Nevertheless, the rapid analysis capability performed with a high accuracy promises that QVG can be an important and valuable tool for the mass analysis of viral samples. The fine-tuning capability of the approach presented here allows the adaptation of the pipeline to a wide range of datasets. Matched with the speed of NGS techniques, QVG could be a tool to track outbreaks by identifying viral strains and checking the within-host diversity of samples.

## Acknowledgments

The pipeline was developed by the bioinformatics group working in the frame of the Complex Health Multidisciplinary Competence Center at the University of Debrecen supported by GINOP-2.3.4-15-2020-00008.

