## Supplemental tables for "Rapid genotyping of viral samples using Illumina short-read sequencing data"

Table S1. Detailed statistics as exported with samtools coverage for the SARS-CoV2 dataset generated for this study.

| sample_id | rname | startpos | endpos | numreads | covbases | coverage | meandepth | meanbaseq | meanmapq |
| --- | --- | --- | --- | --- | --- | --- | --- | --- | --- |
| S23 | MN908947.3 | 1 | 29903 | 194779 | 29652 | 99.16 | 666.72 | 37 | 60 |
| S11 | MN908947.3 | 1 | 29903 | 125861 | 29841 | 99.79 | 439.20 | 37 | 60 |
| S16 | MN908947.3 | 1 | 29903 | 370634 | 29847 | 99.81 | 1148.68 | 36.7 | 60 |
| S24 | MN908947.3 | 1 | 29903 | 491715 | 29869 | 99.88 | 1623.84 | 37 | 60 |
| S10 | MN908947.3 | 1 | 29903 | 822494 | 29865 | 99.87 | 2504.90 | 37 | 60 |
| S5 | MN908947.3 | 1 | 29903 | 55226 | 29449 | 98.48 | 173.48 | 36.9 | 59.9 |
| S12 | MN908947.3 | 1 | 29903 | 454599 | 29889 | 99.95 | 1534.75 | 37 | 60 |
| S9 | MN908947.3 | 1 | 29903 | 1021009 | 29900 | 99.99 | 3301.26 | 37.1 | 60 |
| S14 | MN908947.3 | 1 | 29903 | 1578737 | 29869 | 99.88 | 4621.05 | 37.1 | 60 |
| S15 | MN908947.3 | 1 | 29903 | 168510 | 28308 | 94.66 | 546.672 | 37 | 60 |
| S20 | MN908947.3 | 1 | 29903 | 871661 | 29891 | 99.96 | 2874.43 | 37.1 | 60 |
| S17 | MN908947.3 | 1 | 29903 | 924839 | 29873 | 99.90 | 2878.11 | 37.1 | 59.9 |
| S8 | MN908947.3 | 1 | 29903 | 380861 | 29896 | 99.98 | 1181.52 | 37 | 60 |
| S13 | MN908947.3 | 1 | 29903 | 560779 | 29865 | 99.88 | 1659.33 | 37 | 60 |
| S22 | MN908947.3 | 1 | 29903 | 330740 | 29869 | 99.89 | 1107.72 | 37.1 | 60 |
| S3 | MN908947.3 | 1 | 29903 | 21447 | 28692 | 95.95 | 72.82 | 36.9 | 58.8 |
| S4 | MN908947.3 | 1 | 29903 | 333836 | 29806 | 99.67 | 1094.15 | 36.9 | 60 |
| S2 | MN908947.3 | 1 | 29903 | 441063 | 29872 | 99.90 | 1381.93 | 37 | 60 |
| S18 | MN908947.3 | 1 | 29903 | 735956 | 29870 | 99.89 | 2424.97 | 36.8 | 60 |
| S19 | MN908947.3 | 1 | 29903 | 605008 | 29888 | 99.95 | 2178.96 | 37 | 60 |

Table S2. Detailed statistics as exported with samtools coverage for the publicly available SARS-CoV2 dataset.

| sample_id | rname | startpos | endpos | numreads | covbases | coverage | meandepth | meanbaseq | meanmapq |
| --- | --- | --- | --- | --- | --- | --- | --- | --- | --- |
| SRR14824562 | MN908947.3 | 1 | 29903 | 1770168 | 29903 | 100 | 4134.01 | 36.5 | 60 |
| SRR14824574 | MN908947.3 | 1 | 29903 | 1875745 | 29902 | 99.99 | 4351.20 | 36.6 | 60 |
| SRR14824564 | MN908947.3 | 1 | 29903 | 2453593 | 29896 | 99.97 | 5653.89 | 36.3 | 59.9 |
| SRR14824569 | MN908947.3 | 1 | 29903 | 2452509 | 29898 | 99.98 | 5641.75 | 36.6 | 60 |
| SRR14824568 | MN908947.3 | 1 | 29903 | 2153502 | 29865 | 99.87 | 4966.50 | 36.5 | 60 |
| SRR14824560 | MN908947.3 | 1 | 29903 | 626792 | 29659 | 99.18 | 1460.90 | 36.5 | 60 |
| SRR14824573 | MN908947.3 | 1 | 29903 | 2006313 | 29890 | 99.96 | 4634.16 | 36.6 | 60 |
| SRR14824572 | MN908947.3 | 1 | 29903 | 802442 | 29053 | 97.16 | 1861.43 | 36.6 | 59.9 |
| SRR14824570 | MN908947.3 | 1 | 29903 | 2098095 | 29895 | 99.97 | 4858.03 | 36.4 | 60 |
| SRR14824566 | MN908947.3 | 1 | 29903 | 2088873 | 29877 | 99.91 | 4820.52 | 36.5 | 60 |
| SRR14155385 | MN908947.3 | 1 | 29903 | 2050770 | 29901 | 99.99 | 4584.86 | 36.5 | 59.9 |
| SRR16912539 | MN908947.3 | 1 | 29903 | 114026 | 29903 | 100 | 431.565 | 36.3 | 60 |
| SRR14824563 | MN908947.3 | 1 | 29903 | 2363754 | 29889 | 99.95 | 5446.88 | 36.6 | 60 |
| SRR16741159 | MN908947.3 | 1 | 29903 | 964213 | 29039 | 97.11 | 2230.50 | 36.6 | 59.8 |
| SRR14824561 | MN908947.3 | 1 | 29903 | 2212633 | 29691 | 99.29 | 5113.92 | 36.5 | 60 |
| SRR16912480 | MN908947.3 | 1 | 29903 | 141449 | 29903 | 100 | 529.432 | 36.5 | 60 |
| SRR14824567 | MN908947.3 | 1 | 29903 | 874103 | 29892 | 99.96 | 2030.72 | 36.3 | 60 |
| SRR14824565 | MN908947.3 | 1 | 29903 | 2175466 | 29899 | 99.99 | 5027.86 | 36.6 | 60 |
| SRR17309642 | MN908947.3 | 1 | 29903 | 314900 | 29870 | 99.89 | 488.517 | 37.9 | 60 |
| SRR14155371 | MN908947.3 | 1 | 29903 | 1863425 | 29669 | 99.22 | 4162.81 | 36.5 | 59.9 |

Table S3. Detailed statistics as exported with samtools coverage for the HBV dataset.

| sample_id | rname | startpos | endpos | numreads | covbases | coverage | meandepth | meanbaseq | meanmapq |
| --- | --- | --- | --- | --- | --- | --- | --- | --- | --- |
| SRR12535936 | NC_003977.2 | 1 | 3182 | 249649 | 2752 | 86.49 | 8010.66 | 38 | 59.9 |
| SRR12535937 | NC_003977.2 | 1 | 3182 | 223107 | 2090 | 65.68 | 7357.95 | 37.9 | 59.8 |
| SRR12535946 | NC_003977.2 | 1 | 3182 | 310297 | 3182 | 100 | 9875.27 | 38 | 60 |
| SRR12535938 | NC_003977.2 | 1 | 3182 | 310075 | 3009 | 94.56 | 9586.82 | 37.8 | 59.9 |
| SRR12535947 | NC_003977.2 | 1 | 3182 | 314181 | 3182 | 100 | 9392.32 | 38.1 | 57.8 |

Table S4. Detailed statistics as exported with samtools coverage for the RABV dataset.

| sample_id | rname | startpos | endpos | numreads | covbases | coverage | meandepth | meanbaseq | meanmapq |
| --- | --- | --- | --- | --- | --- | --- | --- | --- | --- |
| SRR12012234 | KT336433.1 | 1 | 11923 | 2675 | 11909 | 99.88 | 15.73 | 36.1 | 59.7 |
| SRR12012235 | KT336433.1 | 1 | 11923 | 79813 | 11916 | 99.94 | 529.40 | 36.1 | 59.9 |
| SRR12012236 | KT336433.1 | 1 | 11923 | 3143 | 11775 | 98.76 | 17.63 | 36 | 59.7 |
| SRR12012237 | KT336433.1 | 1 | 11923 | 144893 | 11923 | 100 | 978.05 | 36 | 59.9 |
| SRR12012238 | KT336433.1 | 1 | 11923 | 106166 | 11915 | 99.93 | 708.35 | 36.1 | 59.9 |
| SRR12012239 | KT336433.1 | 1 | 11923 | 15492 | 11918 | 99.96 | 97.22 | 36.1 | 59.8 |
| SRR12012240 | KT336433.1 | 1 | 11923 | 34641 | 11913 | 99.92 | 222.83 | 36.1 | 59.9 |
| SRR12012241 | KT336433.1 | 1 | 11923 | 34088 | 11912 | 99.91 | 212.80 | 36 | 59.9 |
| SRR12012242 | KT336433.1 | 1 | 11923 | 39417 | 11905 | 99.85 | 246.86 | 36 | 59.9 |
| SRR12012243 | KT336433.1 | 1 | 11923 | 3755 | 11873 | 99.58 | 21.79 | 36 | 59.7 |
| SRR12012244 | KT336433.1 | 1 | 11923 | 1994 | 11685 | 98.00 | 11.206 | 35.9 | 59.5 |
| SRR12012245 | KT336433.1 | 1 | 11923 | 25382 | 11918 | 99.96 | 156.80 | 36.1 | 59.9 |
| SRR12012246 | KT336433.1 | 1 | 11923 | 42129 | 11919 | 99.97 | 276.84 | 36.1 | 59.9 |
| SRR12012247 | KT336433.1 | 1 | 11923 | 13097 | 11905 | 99.85 | 82.51 | 36 | 59.9 |
| SRR12012248 | KT336433.1 | 1 | 11923 | 5038 | 11895 | 99.76 | 29.38 | 36.1 | 59.7 |
| SRR12012249 | KT336433.1 | 1 | 11923 | 10117 | 11911 | 99.90 | 61.76 | 36.1 | 59.8 |
| SRR12012250 | KT336433.1 | 1 | 11923 | 56107 | 11921 | 99.98 | 341.18 | 36 | 59.9 |
| SRR12012251 | KT336433.1 | 1 | 11923 | 18158 | 11912 | 99.91 | 119.40 | 36 | 59.9 |
| SRR12012252 | KT336433.1 | 1 | 11923 | 110577 | 11922 | 99.99 | 671.78 | 36 | 59.6 |
| SRR12012253 | KT336433.1 | 1 | 11923 | 3209 | 11855 | 99.43 | 18.54 | 36 | 59.7 |
| SRR12012254 | KT336433.1 | 1 | 11923 | 17815 | 11914 | 99.92 | 112.33 | 36.1 | 59.9 |
| SRR12012255 | KT336433.1 | 1 | 11923 | 3533 | 11911 | 99.90 | 20.43 | 36.1 | 59.7 |
| SRR12012256 | KT336433.1 | 1 | 11923 | 77218 | 11916 | 99.94 | 497.19 | 36.1 | 59.9 |

Table S5. Detailed statistics as exported with samtools coverage for the FCoV sample.

| sample_id | rname | startpos | endpos | numreads | covbases | coverage | meandepth | meanbaseq | meanmapq |
| --- | --- | --- | --- | --- | --- | --- | --- | --- | --- |
| SRR8352624 | KX722529.1 | 1 | 29174 | 1305 | 27537 | 94.39 | 4.87 | 36.1 | 59.2 |

Table S6. Detailed statistics as exported with samtools coverage for the avian Adenovirus sample.

| sample_id | rname | startpos | endpos | numreads | covbases | coverage | meandepth | meanbaseq | meanmapq |
| --- | --- | --- | --- | --- | --- | --- | --- | --- | --- |
| MT500572 | MG953201.1 | 1 | 45743 | 52014 | 45474 | 99.41 | 121.29 | 33.6 | 58.9 |
